# Long-Horizon associative learning as a unifying framework for statistical learning across scales

**DOI:** 10.1101/2025.03.12.642610

**Authors:** Lucas Benjamin, Ana Fló, Fosca Al Roumi, Ghislaine Dehaene-Lambertz

## Abstract

Sensory inputs are rich with temporal patterns that unfold across multiple timescales. Discovering these regularities is essential for predicting future events and navigating the environment efficiently. Numerous models have been proposed to account for learning at specific temporal scales; however, they are often designed in isolation and rely on narrowly tuned statistical measures, limiting their generalizability to other paradigms. In contrast, humans typically learn without prior knowledge of the underlying structure or the relevant timescale at which regularities occur. Here, we present a unifying account of statistical learning that spans a wide range of temporal dependencies, from adjacent and non-adjacent transitions to complex network structures. This model, Long-Horizon Associative Learning (L-HAL), offers a biologically grounded implementation of the successor representation, or equivalently, the Free Energy Minimization Model. Reanalyzing data from 11 previously published studies, we show that a single neural mechanism, based on graded temporal overlap of associative traces and governed by a single free parameter (β), captures both local statistical regularities and higher-order structural properties. This initial domain-general associative learning process, emerging from the graded structure of associations, may later scaffold to higher-level operations such as grouping, categorization, rule abstraction, and memory formation. Overall, this framework offers a conceptual synthesis that bridges disparate strands of the statistical learning literature and reframes apparent paradigm-specific effects as different expressions of a common underlying computation

## Introduction

Many everyday *inputs, such as speech, music, movements, or actions, unfold as continuous streams of sensory information. These streams are rich, diverse, and temporally structured, embedding complex dependencies across multiple timescales: from immediate transitions between consecutive events *(1)* to more distant regularities linking elements separated by long-range intervals (2–5). Humans are remarkably skilled at discovering these statistical structures. For instance, listeners exposed to continuous speech are sensitive to the likelihood that one syllable follows another, a mechanism proposed to support early word learning in infants (1, 6). They also detect non-adjacent linguistic relationships, such as “is”…”ing” regardless of the number of intervening syllables (4, 7), and even infer higher-order structure in network-like sequences (3, 8–12): when stimuli are drawn from distinct graph-based clusters, participants detect transitions between clusters even when adjacent transition probabilities remain constant.

Although some attempts have been made to aggregate adjacent and non-adjacent dependency learning under a common associative framework (13), much of the statistical and network learning literatures still relies on distinct models tailored to specific experimental designs. These models can be broadly categorized by the temporal scale they target: local (transitions between adjacent elements), intermediate (non-adjacent dependencies), and high-order (network-level structure) (***Fig. 1***). This multiplicity implicitly assumes that participants engage in numerous parallel, scale-specific computations, a fragmentation that poses serious memory and efficiency challenges, especially for infants, whose cognitive resources, even if limited, support impressive learning capacities (1, 4, 14, 15).

**Fig. 1.**
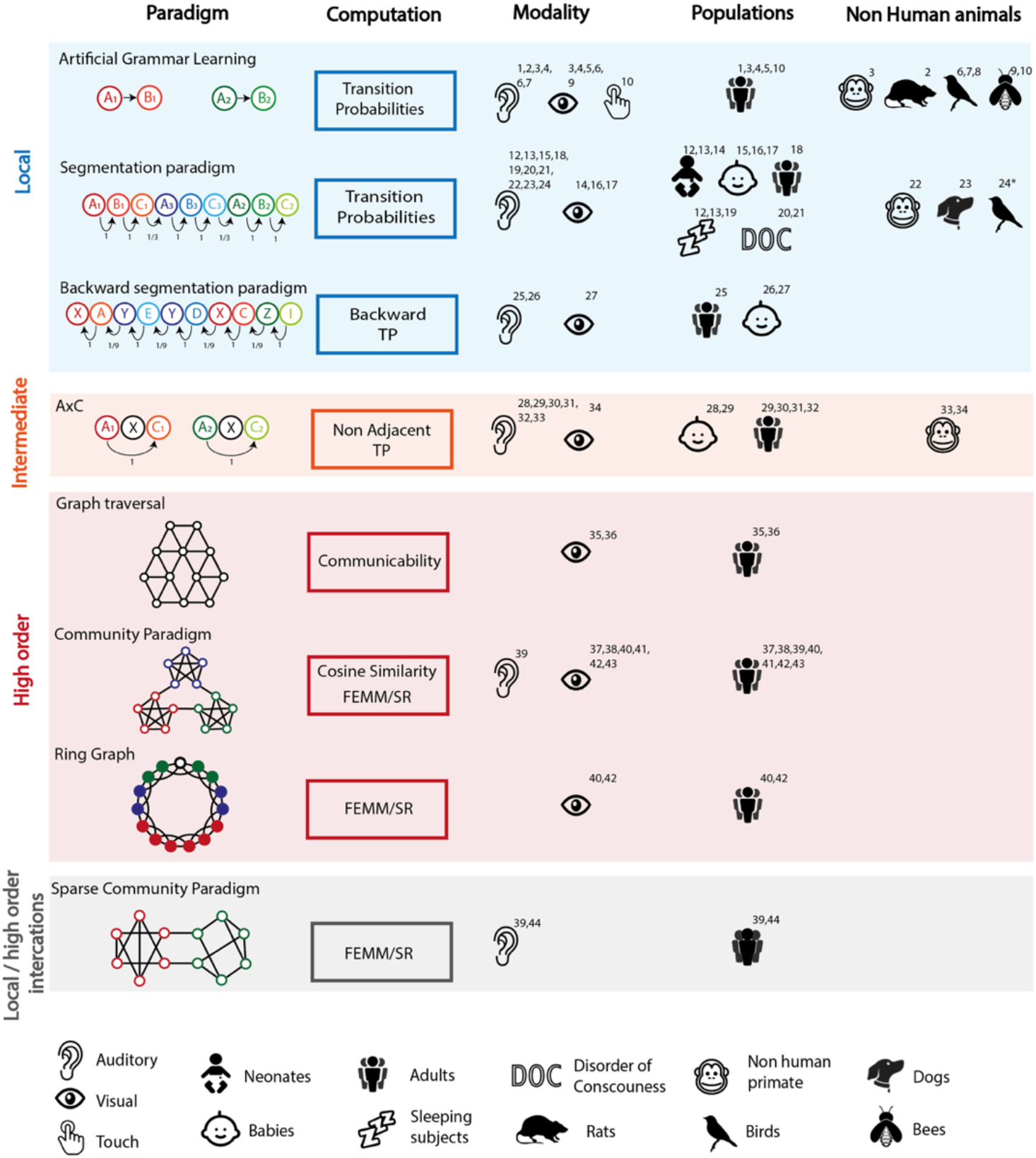
Overview of statistical learning paradigms categorized by the scale of temporal dependencies: local, intermediate, and high-order. The figure highlights some studies demonstrating learning at different scales of statistical structure, across various sensory modalities and participating populations. It should be noted that this does not necessarily represent the ability of populations to learn or not learn each paradigm, but rather the current state of our knowledge on this subject and on what has been formally tested. **1**. *Harris* (6) **2**. *Toro & Trobalon* (62) **3**. *Milne et al* (30) **4**. *Conway & Christiansen* (2006) **5**. *Reber* (1967) **6**. *Santolin et al* (63) **7**. *James et al* (64) **8**. *Menhart et al* (65) **9**. *Avarguès-Weber et al* (66) **10**. *Comway & Christiansen* (67) **11**. *Nicholls et al* (68) **12**. *Fló et al* (23, 24, 28) **13**. *Benjamin et al* (14) **14**. *Bulf et al* (21) **15**. *Saffran et al* (1) **16**. *Fiser & Aslin* (22) **17**. *Kirkham et al* (25) **18**. *Saffran et al* (1, 26) **19**. *Batterink & Zhang* (19) **20**. *Xu et al* (27) **21**. *Benjamin et al* (20). **22**. *Hauser et al* (69) **23**. *Boros et al* (70) **24**. *Takahasi* (71) **25**. *Perruchet and Desaulty* (72) **26**. *Pelucchi et al* (73) **27**. *Tummelsthammer et al* (74) **28**. *Buiatti et al* (31) **29**. *Gómez et al* (7) **30**. *Peña et al* (4) **31**. *Endress* (75) **32**. *Newport & Aslin* (15) **33**. *Newport et al* (76) **34**. *Sonnweber et al* (77) **35**. *Garvert et al* (3) **36**. *Mark et al* (11) **37**. *Schapiro et al* (5, 32) **38**. *Karuza et al* (35, 36) **39**. *Benjamin et al* (8) **40**. *Lynn et al* (9) **41**. *Ren et al* (39) **42**. *Stiso et al* (12) **43**. *Kakaei et al* (78) **44**. *Benjamin et a*l (18)

To reconcile learning across timescales within a unified framework, we suggest that both short- and long-range associations arise from a single neural mechanism grounded in Hebbian principles, whereby learning results from overlapping neural representations of successive elements (16). Because neural traces decay gradually, successive elements overlap in time, enabling associations not only between consecutive events but also between temporally distant events, creating a “long-horizon” associative field. This mechanism is both parsimonious and biologically plausible, relying on sustained patterns of neural activity that could, in principle, be implemented with a limited number of neurons. It can also be formalized mathematically, yielding parameters that can be directly linked to measurable neural properties.

The Free Energy Minimization Model (FEMM (9)) and the Successor Representation (SR (17)) model provide formal instantiations of this long-horizon associative learning (L-HAL). Initially proposed to explain network learning in humans (FEMM) and cognitive map formation for spatial navigation (SR), the two models are mathematically equivalent despite their distinct conceptual origins. Both describe associative strength as a linear combination of the transition probability matrix for adjacent elements (A_1_) and higher temporal orders (A_Δt^*^), each weighted by an exponential decay with Δt^†^. A single parameter, β, governs this decay: when β → ∞, only adjacent transitions contribute; when β → 0, all elements become equally associated regardless of distance. Thus, estimating β directly specifies the associative timescale. The model is agnostic to the nature or granularity of the elements, phonemes, syllables, tones, or other primitives. It describes an early, domain-general associative process whose output may scaffold higher-level operations such as grouping, categorization, rule abstraction, and memory formation. We previously tested the plausibility of such long-horizon associations using an auditory “sparse community” paradigm in which tone sequences were governed by both local and high-order dependencies. Tones were drawn from a network organized in two clusters, so that transitions occurred either within or between clusters while adjacent transition probabilities were held constant. Participants later judged within-cluster transitions as more familiar, even when they had never been presented; whereas between-cluster transitions were judged less familiar, despite being presented as often as the within-cluster ones (8) (see ***Fig. 1***, Local/High-order interaction). Participants were thus sensitive to the higher-order structure of the network, implicitly completing missing transitions consistent with that structure while rejecting those that violated it. A subsequent MEG study revealed neural signatures consistent with decaying-overlap dynamics(18), supporting the idea that even high-order structure learning and generalization can emerge from a very simple associative rule.

Given its general formulation, the L-HAL framework should capture temporal associations at all scales, from adjacent to long-range dependencies. We therefore challenged the model with a broad set of published datasets spanning local, non-adjacent, and network-level learning. The β parameter may vary across domains, developmental stages, or cognitive states. However, we might expect it to fall within a restricted range that captures the main signatures of human learning, thereby offering a unified, parsimonious account of statistical learning across scales. Thus, we provide an explicit comparison of the data with model predictions using a unique β value previously optimized (8, 9), and evaluate domain validity from predictions obtained across a broad range of β values.

## Model description

Our analyses drew on publicly available datasets encompassing a wide range of behavioral and neurophysiological measures, including familiarity rankings, reaction times in healthy adults, looking preferences in infants, ERP and fMRI. Collectively, these datasets cover the full range of temporal scales depicted in Fig. 1. We qualitatively compared the experimental results with the model’s predictions and quantitatively assessed the goodness of fit when sufficient data were available.

### Formal description of the model

Given the formal equivalence between FEMM and SR (see discussion) and for the sake of consistency with our previous work, we anchored the description in the FEMM formalism. This model captures a trade-off between short and long-distance associations: instead of linking only consecutive elements (i and i–1), it also binds each element i to earlier ones (i–2, i–3, etc.) with a probability that decreases exponentially with temporal distance, following a Boltzmann distribution (See SI Appendix).

The estimated model can then be written as:

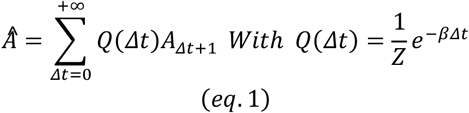

***A***_Δ*t*_: Non-adjacent transition probability matrix of order Δ***t*** between elements of the sequence,

**β**: Generalization factor varying from 0 (no generalization) to 1 (uniform generalization),

**Z:** Constant value *Z*=1/(1−*e*^− β^)

Δ ***t***: Distance between items of the sequence.

For sequences following a Markovian rule - the current element is only dependent on the previous one - ***A***_Δ ***t***_=***A***^Δ***t***^. Eq. 1 can be rewritten:

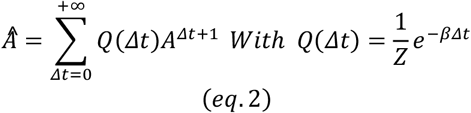

Which simplifies as (See SI Appendix):

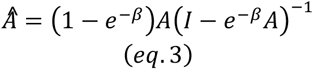

**I**: Identity matrix (same size as A)

### Computation of the model for each paradigm

We carefully curated landmark studies spanning multiple scales of statistical learning. They mostly shared a common structure: participants were exposed to sequences of stimuli during training, then tested on congruent and incongruent items in a post-learning phase. Learning was inferred from differences in familiarity ratings or deviance responses between these item types.

We simulated the associative strength between elements from the training material, computed the predicted familiarity for each test condition, and compared these predictions with published empirical results. To maintain parsimony and avoid paradigm-specific overfitting, we first fixed β to be 0.06, the value proposed by Lynn et all(9) and that we also found in our previous work(8). As a second step, for each study, we identified the range of β values yielding the best fit, exploring values from 10^−4^ to 10^2^ to determine the optimal range explaining performance across paradigms.

The following approach was used:

– For Markovian sequences, we computed the analytical association matrix using eq 3.
– For AXC and artificial grammar paradigms, where A is insufficient to fully describe the training material, we estimated FEMM associations directly from the corpus of training sequences. This was done by computing and summing (with an exponential decay factor) all TP orders within each training sequence (eq. 1). The final 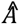 matrix was obtained by averaging the estimated association matrix across all training sequences experienced by participants.

This method estimated the optimal learning after convergence, ignoring possible local imbalance or curriculum effects (see later in the discussion for a model including learning dynamics). Once the participants’ theoretical optimal mental model was derived from the training material, we computed the predicted familiarity for each test condition:

– For a pair (*ab)*, the estimated familiarity is 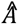 (*ab*). This is the case for all network paradigms (Fig. 3 E,F,G&I)
– For longer test sequences, we computed a weighted sum of all pairwise associations between elements. For instance, the familiarity of a triplet *abc* was estimated as: 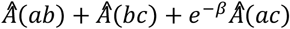. This procedure was applied to all paradigms involving test sequences with more than two elements (Fig. 3A,B,C&D). In the artificial grammar paradigm, the state “end” was treated as an item, following the original study.
– In the specific case of Lynn et al(9), we followed their original method to compare two types of networks: Community and Lattice structures, by computing the average FEMM familiarity across all transitions within each network.

We obtained quantitative predictions of model familiarity for each experimental condition and compared them with published experimental measures.

## Results

All the results are presented in Figure 3.

### Local scale

#### Segmentation paradigms

Sensitivity to adjacent transition probabilities has long been recognized as a core mechanism of statistical learning, most notably demonstrated in the seminal speech segmentation study in 8-month-old infants*(1)*. In such paradigms (1, 14, 19–27), participants hear continuous streams of syllables (or other auditory, visual, or haptic stimuli) containing pseudo-words (ABC). Within words, transition probabilities are high (TP=1), but drop to 1/3 across word boundaries. After brief exposure, participants are shown to have successfully extracted words (A_i_B_i_C_i_) as they distinguish them from part-words (B_i_C_i_A_k_) straddling a word boundary, through differences in looking time, familiarity rating, chunk recall accuracy, or EEG responses (14, 20, 24, 26–29).

#### Comparison with the model

FEMM simulations reproduced these effects: for β = 0.06, word familiarity exceeded part-word familiarity by 19% (Fig 3A). As adjacent transition probability is the first order and dominant component of the FEMM, it is not surprising that a TP difference induced a FEMM difference and that larger β values further amplify the word vs part-word difference (Fig 3A).

### Grammar learning paradigms

Milne et al.’s grammar-learning study (30) offered an ideal test case, combining simplicity with fully reported training and test sequences. Human participants and macaques were exposed to 8 different training sequences, then judged whether new five-item sequences followed the same grammar. Their judgments correlated with local transition probabilities in both visual and auditory modalities for both humans and macaques.

#### Comparison with the model

FEMM predictions, computed from the training set, closely matched participants’ behavior (Visual R=-0.69 p<0.05, Auditory R=-0.74 ps<0.01 for β =0.06 – Fig. 3B), indicating that the model accurately captured observed learning patterns. This was not the case for macaques: only higher β values provided a good fit to their data, suggesting that monkeys primarily encoded first-order transition probabilities, neglecting longer-distance associations (See SI Appendix Fig. S2).

### Intermediate scale

#### AXC paradigm

At the intermediate level, sensitivity to non-adjacent dependencies has been proposed to underlie grammatical learning, such as tense agreement in English (“is … ing”, “has … ed”). Peña et al(4) showed that adults can learn non-adjacent transition probabilities of the form P(C|XA), where the syllable A predicts C despite an intervening X syllable when AXC triplets were separated by subliminal pauses. In that experiment, participants later preferred test items preserving the A-C dependency (Rule Word AX’C where X’ is novel) over Part-Words (XCiAk). Gómez (7) further found that learning improved as the variability of possible Xs increased (2, 6, 12, or 24 syllables), a result that cannot be explained by learning the constant A-C dependency alone.

#### Comparison with the model

We estimated FEMM familiarity scores for Words, Part-words, and Rule-Words in the classical case with three possible intervening Xs (31) and training on isolated words. Following the authors’ interpretation that pauses segmented the stream, we computed the model on each triplet (Fig. 1 in (4), Fig. 3C). For β = 0.06, the model predicted Words familiarity (A_i_XC_i_) > Rule-Words familiarity (A_i_X’C_i_) > Part-Words familiarity (XC1A2), closely matching behavioral data(4, 15, 31). Consistent with empirical findings(4), removing pauses during training (i.e., using a continuous stream) abolished the familiarity advantage for Rule-Words over Part-Words.

Varying the number of possible Xs (2–29) replicated Gómez’s result: the Word > non-Word familiarity difference increased with X variability (*8 - Fig3D*). This pattern reflects FEMM’s trade-off between local and distant associations: When X variability grows, adjacent transitions (AX, XC) weaken while the A–C link remains constant, biasing predictions toward the non-adjacent dependency. Finally, unlike in local paradigms, the best-fitting β range for intermediate-scale learning shifts toward lower values, but still displays a good overlap with the local paradigm range of validity (Fig. 3C–D).

### Higher order scale

#### Garvet et al’s network paradigm

At a higher-order scale, sequence structure is often described using network formalisms. Garvert et al. (3) showed that hippocampal–entorhinal fMRI activity during navigation of a 12-node network correlated with Communicability, graph-based measure of distance between nodes. Although conceptually similar to FEMM or SR, Communicability penalizes long-range dependencies more strongly.

#### Comparison with the model

We computed FEMM familiarity estimates for each network transition and re-analyzed the average activation within the hippocampal–entorhinal cluster identified by Garvert et al. Mean activation per transition was significantly correlated with FEMM predictions (R=-0.29, p<0.05 for β=0.06) (Fig. 3E), supporting the model’s relevance for explaining brain responses to structured sequences.

### Community network

Schapiro et al.(5) introduced the community network paradigm to demonstrate humans’ sensitivity to network structure, even in the absence of informative local statistics. The network consists of three communities—groups of nodes highly interconnected internally but weakly connected to other groups—where each node has exactly four neighbors, thus a uniform probability of 0.25. Despite this flat local structure, participants reliably distinguished transitions within communities from those between communities (5, 8, 9, 18, 32–36). Lynn et al.(9) further contrasted this community network with a lattice network of identical size and average connectivity and found that participants trained on a lattice structure exhibited lower familiarity (slower reaction times) compared to those trained on a community structure.

#### Comparison with the model

Originally proposed to explain these effects, FEMM accurately predicted the observed drop in familiarity at community boundaries (9) (Fig. 3F) and reproduced the higher overall familiarity for community versus lattice networks. Indeed, average FEMM familiarity across all transitions was higher for the community network than for the lattice network, consistent with participants’ faster reaction times when learning the community structure (Fig. 3H).

### Ring graph

Lynn et al. also investigated a different network topology: a 15-node ring-graph in which each node connects to four neighbors (+1/+2/-1/-2). Thus, nodes are separated by one, two or three transitions. Measuring the familiarity of each pair of this network using reaction times revealed a graded preference for directly connected pairs (distance 1), followed by unconnected pairs at distance 2, and then distance 3.

#### Comparison with the model

FEMM reproduced this hierarchy of familiarity, as shown by Lynn and colleagues(10) (Fig. 3G), with predicted familiarity decreasing systematically with increasing graph distance.

### Local and high-order interaction

#### Sparse and high sparse community networks

In previous studies, we introduced the sparse community design(8, 18), in which one (sparse) or two (high sparse) possible transitions per node inside each community were removed. It created a 2-by-2 design used during test where a transition between two sounds could be Familiar or New transitions (heard vs. unheard during training) and occurred Within or Between communities (the two sounds were from the same community or not). Participants systematically “completed’’ the missing within-community transitions—judging them as familiar despite never hearing them— indicating they reconstructed the latent structure. Conversely, between-community transitions were judged less familiar than within-community ones, even though both occurred equally often.

#### Comparison with the model

To explain this behavior, we systematically compared leading models from the statistical and network learning literature, including transition probabilities, non-adjacent transition probabilities, n-grams, communicability… FEMM provided the best fit(8, 18) (Fig. 3I). Remarkably, across the full, sparse, and high-sparse community designs, only a narrow range of β values fitted the data (Fig. 3I), making these paradigms powerful probes for pinpointing the associative timescale.

## Discussion

How to efficiently extract multiple scales of structure from temporal input? This question has been of great interest to the scientific community for a long time, both for human cognition and artificial neural networks, and has led to the introduction of many computational innovations in artificial learning agents, including the emergence of recurrent connections in artificial networks (37), or memory traces learning methods for spiking networks (38). The framework for statistical learning presented in this study seeks to connect formal mathematical and biological descriptions, explaining how humans learn conditional probabilities across different temporal structures. Indeed, its dynamics can be instantiated without recurrent connections, by simple physiologically plausible mechanisms such as sustained neural activity. When items’ representations persist over time, their temporal overlap lets the brain form associations across various time-scales. This mechanism enables the simultaneous learning of a wide range of conditional probabilities within a sequence. These persistent neural traces are assumed by the models to decay exponentially (Fig. 2B), a hypothesis supported by recent empirical evidence (20, 21).

**Fig. 2.**
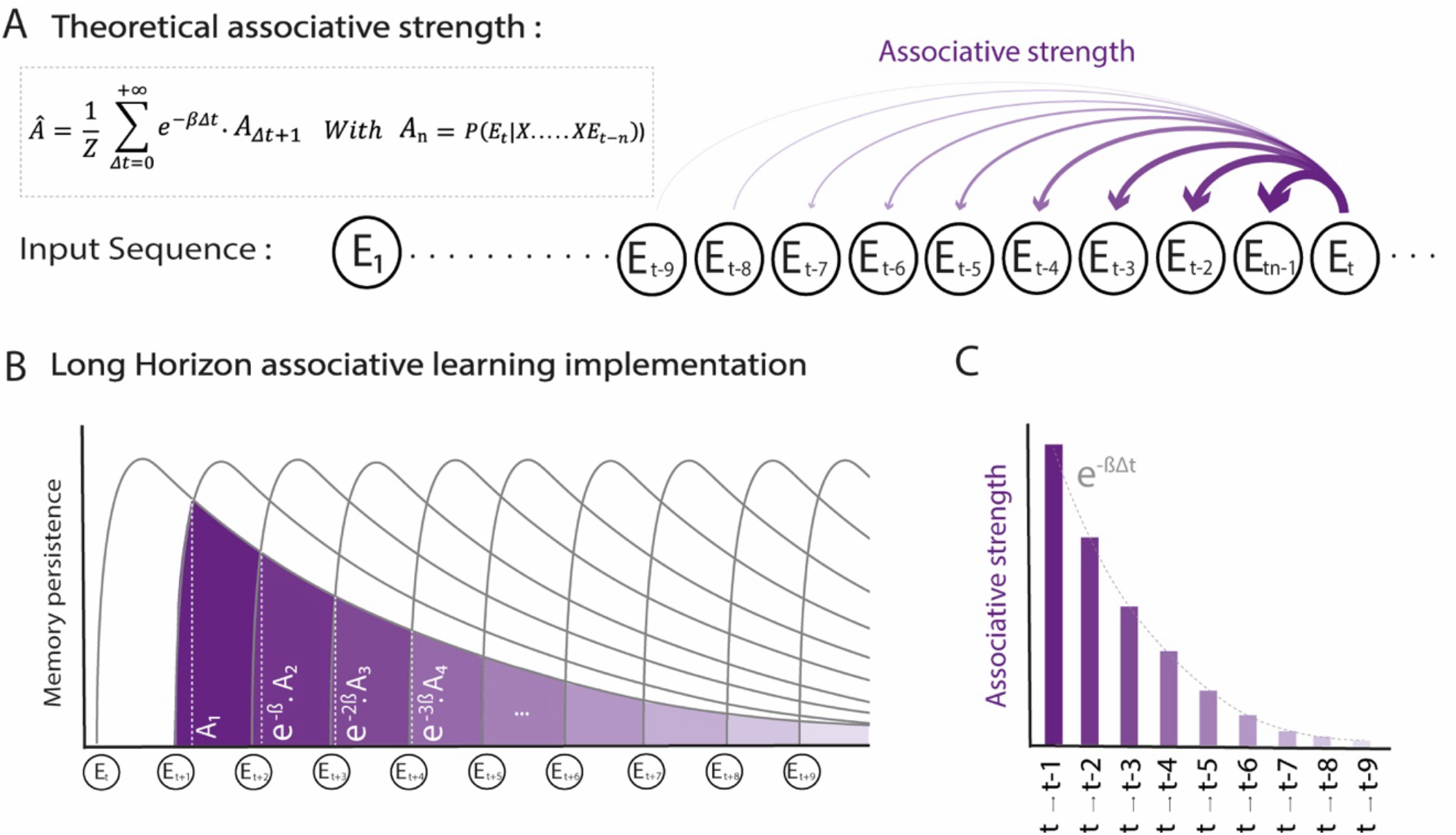
Description of the FEMM and its Long-Horizon Associative learning implementation. **A**. Formal equation of the FEMM, used to compute the association matrix (Â) between all pairs of elements in a sequence. Â reflects the cumulative contribution of adjacent and non-adjacent transition probabilities across all temporal distances, weighted by an exponential decreasing function. The diagram on the right illustrates the associative strength between a current element Et and its predecessors. It shows strong association with the immediately preceding element (Et-1, adjacent TP) and weaker associations with earlier elements (Et-2, Et-3….; non-adjacent TP) consistent with the exponential decay. **B**. Graphical representation of the FEMM’s implementation through a Long-Horizon Associative Learning strategy. If the mental representation of each element decays exponentially over time, the overlap between successive representations allows for Hebbian associations to form between both adjacent and non-adjacent elements, following a penalty computed with FEMM. **C**. Schematic depiction of the exponentially decaying associative weights between the current element and its predecessors, illustrating connections to both consecutive (t -> t-1) and nonconsecutive (t –> t-2; t -> t-3; …) elements of the sequence

**Fig. 3.**
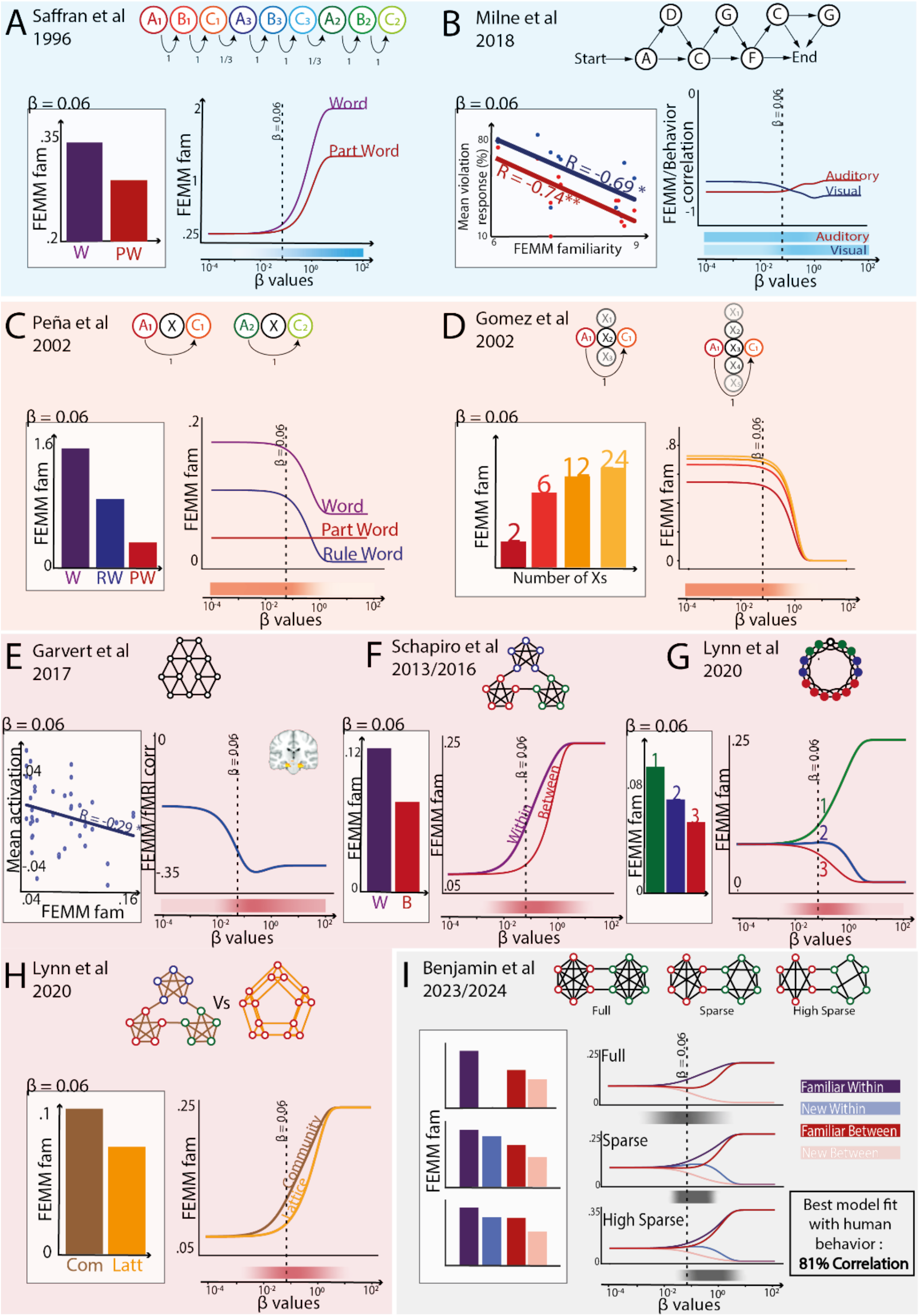
Results from the analysis of the statistical learning paradigms across different temporal scales using the Free-Energy Minimization Model (FEMM). In this figure, the left panels in white display the model predictions for a previously optimized β value (β=0.06), and the right panels the model predictions for β values between 10^−4^ and 10^2^. Dotted lines show β=0.06, and shaded bars below illustrate the β range for which the model accurately predicts the available human data. **(A)** In word segmentation tasks using familiarization with continuous speech streams, participants are subsequently more familiar with embedded “Words” than with “Part-Words” that span boundaries. For this local paradigm, higher β values better predict the Word vs Part-Word differences. **(B)** Artificial grammar learning paradigms target local transition probability learning. FEMM familiarity values significantly correlate with mean violation responses reported by Milne et al. (30) in both auditory (red) and visual (blue) modalities. Modeling of macaques’ performances is reported in SOM. **(C)** At the intermediate scale, sensitivity to non-adjacent dependencies has been demonstrated using AXC paradigms. The reported preference for RuleWord over PartWord is also correctly accounted by FEMM. **(D)** Increasing the number of X elements in the AXC paradigm affects participants’ performances in a Word vs NonWord discrimination task, despite the non-adjacent transition probability P(C|XA) = 1 remaining constant across conditions. FEMM estimates (left bar-plot) for varying numbers of intervening X elements (ranging from 2 to 29) mirror the behavioral results reported by Gómez et al. (7) (right bar-plot), showing that word preference increases with the number of possible intervening syllables. **(E)** Using fMRI, Garvert et al. (3) showed that when exposed to a sequence deriving from a network structure, a cluster in the entorhinal cortex significantly correlated with Communicability. FEMM also shows a significant correlation with the average activation within this cluster. **(F)** In the community paradigm (5), participants distinguished transitions within vs. between clusters despite identical local transition probabilities. FEMM reproduces this distinction as shown in the bar plot comparing the familiarity of within and between transitions. **(G)** In a ring graph paradigm, Lynn et al. (9) showed graded familiarity judgments based on graph distance (2 vs. 3 steps apart). FEMM accurately predicts these gradations. Distance 1 corresponds to the connection between white and green nodes, Distance 2 to white and blue nodes, and Distance 3 to white and red nodes. **(H)** Lynn et al(9) also showed that learning a community structure was easier (faster reaction time) than a lattice structure. FEMM predicts this effect, with higher average familiarity for community (brown) than lattice (orange) graphs. **(I)** In a modified version of the community paradigm (sparse and highly sparse designs), we removed some within-community transitions to be able to test learning of local and high-order structures. FEMM predicted generalization within-community, assigning higher familiarity to non-existent within-community transitions than to familiar between-community ones. Behavioral and MEG results confirmed this prediction. We showed that FEMM significantly outperformed the other statistical models proposed in the literature

Our goal was to show that this single associative principle can account for the diversity of findings reported in the statistical learning literature, across multiple temporal scales. While the Free-Energy Minimization Model (FEMM) provided the original mathematical formalization, it was originally developed in the predictive coding framework. Here, we adopt the more neutral term Long-Horizon Associative Learning (LHAL) to emphasize that the same formal mechanism can arise passively from overlapping neural traces, without invoking explicit prediction or error correction. The model accurately reproduces experimental data while generalizing across paradigms that have traditionally required distinct, domain-specific models. More broadly, effects interpreted as abstract rule learning (35, 36)—such as nonadjacent (AXC) dependencies or network completion—may instead emerge naturally from long associative dynamics. It also bridges two domains often studied independently: statistical learning and network learning. While network learning is traditionally viewed as abstract structure learning requiring explicit mental maps(39, 40), our results suggest that these high-order effects can emerge naturally from the same implicit associative dynamics seen in statistical learning. Finally, the model’s minimal architecture makes it biologically plausible: learning across timescales reduces to a single Hebbian process governed by gradually decaying memory traces.

### A Single Free Parameter to Capture Integration Across Statistical Scales

LHAL explains how associations extend across time through a single parameter, β, which tunes the balance between precision and generalization. High β values restrict learning to local transitions, while low values promote integration across distant elements. Estimating β across species or ages could thus reveal differences in associative timescales and generalization capacity. Cross-species data support this interpretation: in Milne et al. (30), macaques’ behavior was best captured by high β values, indicating reliance on first-order associations (See SI Appendix Fig. S2). In human infants, β may start high—reflecting immature neurons’ limited capacity for sustained firing—favoring short-range associations, consistent with evidence of local transition learning in sleeping neonates. Implicit learning paradigms combined with EEG recordings could be used to track how β evolves with age. As such, β may provide a testable functional link between regional brain maturation and changes in performance.

Note, however, that the measured β is often not derived directly from the output of the LHAL mechanism, but rather from the behavioral outcomes of the decision and response processes. For instance, in an active and explicit behavioral version of the sparse community paradigm, we estimated an average β of 0.06. In contrast, using the same paradigm in an implicit and passive MEG study yielded a higher estimate of β=0.5. While this difference may partly result from variations in measurement sensitivity, it may also reflect the impact of active engagement. The effort to memorize, by recruiting stronger top-down processes, could contribute to higher maintenance of remote elements (longer traces = lower β – Fig 2A&B). Given ongoing debates about the role of attention in statistical learning(20, 27, 41–46), careful estimation of β across different paradigms might help clarify the exact role of attention in learning.

Finally, significant individual differences might also modulate the β value. Although challenging to measure at the individual level, Lynn and colleagues(9) showed that an individually fitted β parameter correlated with participants’ memory capacity in an n-back task, providing initial evidence that individuals may differ in their generalization/accuracy trade-off, or possibly reflecting attentional engagement in both tasks. We still lack intra-subject test-retest reliability of this parameter to investigate this question. Given the focus on measurement reliability in the recent literature(29), a specific study on the consistency of individual β parameters across statistical learning tasks is highly necessary.

### Biological implementation

We proposed Long-Horizon Associative learning as a biologically grounded implementation of FEMM or SR. Such an implementation predicts traces of past elements slowly decaying with time. In a previous MEG experiment, we found evidence consistent with this: neural representations of tones decayed exponentially over time (or number of elements – see limitation paragraph), with overlapping activity across up to eight successive tones (Fig. 2B & C). Moreover, MEG signals reflected the probability of upcoming tones based on this decay pattern, capturing associations at multiple orders of transition probabilities(18). Recently, Kahn and colleagues(34) found learning properties compatible with trace like implementation of SR, rather than an alternative encoding with element reactivations. This further supports Long-Horizon Associative learning as a good candidate to implement FEMM or SR in implicit sequence learning tasks.

### Model extension to spatial associations

The principles underlying associations between temporally ordered events (a single dimension) described here can, theoretically, be extended to any space, regardless of the axes or the number of dimensions. In particular, for spatial patterns, the association strength (Fig 2B) can be defined by replacing Δt by Δdistance, with Δdistance = max(|Δx| + |Δy|), or any other 2D distance metric (See SI Appendix). While this formulation provides a computational analogue of FEMM for spatial patterns, the equivalence does not hold at the level of neural implementation. Implementing such two-dimensional long-distance associations would likely require mechanisms beyond simple sustained neural activity (e.g., the serial allocation of attention to limited spatial regions of a scene, noisy grid-cell, etc). Characterizing these mechanisms, however, lies beyond the scope of the present manuscript.

Related ideas have emerged in the reinforcement learning and spatial navigation literatures. In these domains, hippocampal place cells have been proposed to represent maps of probabilistic future states and rewards by encoding the successor representation rather than simple positional cognitive maps(17, 47, 48). Because future states may encompass multiple steps ahead, the underlying mathematical problem is analogous to integrating transition probabilities across different temporal orders. The successor representation (SR) provides a formal solution to this problem. SR is defined as a weighted sum of probabilistic future states: SR = ∑_Δ***t***_ γ^Δ*t*^ ***A***^Δ*t*^, where A is the transition probability matrix between successive states and γ^Δ*t*^(with 0 < γ < 1) is a discount factor. Although FEMM and SR emerged from different research contexts (statistical learning vs. cognitive map), they are mathematically equivalent under the transformation γ = *e*^−β^. This convergence suggests that similar computational principles may underlie sequential statistical learning, spatial pattern recognition, and spatial navigation, albeit with distinct neural implementations.

### Model limitations and further directions

Our goal here was to highlight the power of low-level associative mechanisms and their potential role in supporting a wide range of statistical learning phenomena. However, such mechanisms are unlikely to explain all aspects of human sequence learning. In many cases, additional processes may build upon this initial layer, whose output might be further integrated into higher-level structures (49) —such as syllables into words —on which long-horizon associative learning may operate again. Statistical learning in speech is also shaped by prosodic boundaries (14, 50, 51), suggesting that associative learning is constrained by the structural domain in which it occurs.

Because L-HAL formalizes a graded associative process based on overlapping neural traces, it inherently supports probabilistic generalizations that extend beyond the observed input. It enables extrapolation to unobserved but statistically compatible combinations, such as “phantom words”(52) or graph completion in the sparse community paradigm (8, 18). However, it is radically different from rule-based learning (53–58), which involves explicit abstraction: the input is no longer encoded as sensory exemplars but instead represented in terms of labels. For instance, two alternating tones of 800 and 1000 Hz may be abstracted as an ABAB pattern, and generalized to any alternating stimuli, tones, or visual stimuli (58). This symbolic form of generalization is not accessible to L-HAL and likely emerges at a later processing stage.

Further research is needed to clarify how these forms of learning interact and whether rule-based learning might emerge as a strategy to reduce the cognitive cost of maintaining long-range associations. It remains also to be determined whether the decay in associative strength is governed primarily by time or by the number of intervening elements (as implemented here). Fig. 3 showed that for most paradigms, the model predictions are not drastically different with a 10-fold β factor. Within a range of medium-paced sequences, considering time or number of elements would yield similar results for short elements like syllables or tones.

In this paper, the model is initially described using a theoretical transition matrix to compute the familiarity estimate. However, this approach overlooks the effects of incomplete learning during sequence exposure, as well as curriculum effects such as recency or primacy bias for instance (59–61). The model can be readily extended to address these limitations by replacing the absolute familiarity estimate with a dynamic one, updated after the presentation of each element *i*. Formally, *eq. 1* is replaced by:

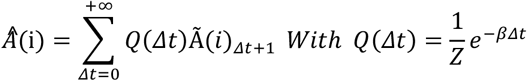

Where Â (i) is the model estimation after the presentation of element i in the sequence and Ã (i) the estimated transition probability conditioned on the sequence up to element i. This matrix reflects the actual sequence experienced by subjects and can incorporate biases (for instance, recency bias can be implemented by restricting Ã (i) to recently observed items). In this formulation, Long-Horizon associative learning becomes a dynamic framework that captures both online learning during sequence presentation and the amount of exposure required to learn each sequence, as well as curriculum-related effects. Such a dynamic formulation could help account for individual learning trajectories rather than just the final result of efficient learning as in the current study.

## Acknowledgments

This work received funding from the European Research Council (ERC) under the European Union’s Horizon 2020 research and innovation program (grant agreement No. 695710 and 101142651) and from the French government managed by the Agence Nationale de la Recherche as part of the France 2030 program, under reference ANR-23-IAHU-0010. LB thanks Les Treilles Foundation for its support.

The authors used an AI-assisted language tool to improve the clarity and grammar of the manuscript. All scientific content and interpretations are the sole responsibility of the authors.

## Availability statement

All codes for this study are available at https://osf.io/g65d7

## Footnotes

† Δt corresponds to the distance between items of the sequence and not time per se.

## Notes

### Competing Interest Statement

The authors have declared no competing interest.

### Summary of Updates

Overall re-writting, ading discussion points and limitations, more comprehesnive bibliography

## References

1. J. R. Saffran, R. N. Aslin, E. L. Newport, Statistical Learning by 8-Month-Old Infants. Science 274, 1926–1928 (1996).

2. A. Basirat, S. Dehaene, G. Dehaene-Lambertz, A hierarchy of cortical responses to sequence violations in threemonth-old infants. Cognition 132, 137–150 (2014).

3. M. M. Garvert, R. J. Dolan, T. E. Behrens, A map of abstract relational knowledge in the human hippocampal– entorhinal cortex. eLife 6, 1–20 (2017).

4. M. Peña, L. L. Bonatti, M. Nespor, J. Mehler, Signal-Driven Computations in Speech Processing. Science 298, 604– 607 (2002).

5. A. C. Schapiro, T. T. Rogers, N. I. Cordova, N. B. Turk-Browne, M. M. Botvinick, Neural representations of events arise from temporal community structure. Nat. Neurosci. 16, 486–492 (2013).

6. Z. S. Harris, From Phoneme to Morpheme. Language 31, 190 (1955).

7. R. L. Gómez, Variability and Detection of Invariant Structure. Psychol. Sci. 13, 431–436 (2002).

8. L. Benjamin, A. Fló, F. Al Roumi, G. Dehaene-Lambertz, Humans parsimoniously represent auditory sequences by pruning and completing the underlying network structure. eLife 12, e86430 (2023).

9. C. W. Lynn, A. E. Kahn, N. Nyema, D. S. Bassett, Abstract representations of events arise from mental errors in learning and memory. Nat. Commun. 11, 2313 (2020).

10. C. W. Lynn, D. S. Bassett, How humans learn and represent networks. Proc. Natl. Acad. Sci. 117, 29407–29415 (2020).

11. S. Mark, R. Moran, T. Parr, S. W. Kennerley, T. E. J. Behrens, Transferring structural knowledge across cognitive maps in humans and models. Nat. Commun. 11, 1–12 (2020).

12. J. Stiso, et al., Neurophysiological Evidence for Cognitive Map Formation during Sequence Learning. eneuro 9, ENEURO.0361-21.2022 (2022).

13. A. D. Endress, S. P. Johnson, When forgetting fosters learning: A neural network model for statistical learning. Cognition 213, 104621 (2021).

14. L. Benjamin, et al., Tracking transitional probabilities and segmenting auditory sequences are dissociable processes in adults and neonates. Dev. Sci. 26, e13300 (2022).

15. E. L. Newport, R. N. Aslin, Learning at a distance I. Statistical learning of non-adjacent dependencies. Cognit. Psychol. 48, 127–162 (2004).

16. D. O. Hebb, The Organization of Behavior (Lawrence Erlbaum, 2002).

17. P. Dayan, Improving Generalization for Temporal Difference Learning: The Successor Representation. Neural Comput. 5, 613–624 (1993).

18. L. Benjamin, M. Sablé-Meyer, A. Fló, G. Dehaene-Lambertz, F. Al Roumi, Long-Horizon Associative Learning Explains Human Sensitivity to Statistical and Network Structures in Auditory Sequences. J. Neurosci. 44, e1369232024 (2024).

19. L. J. Batterink, S. Zhang, Simple statistical regularities presented during sleep are detected but not retained. Neuropsychologia 164, 108106 (2022).

20. L. Benjamin, et al., The role of conscious attention in auditory statistical learning: Evidence from patients with impaired consciousness. iScience 28, 111591 (2025).

21. H. Bulf, S. P. Johnson, E. Valenza, Visual statistical learning in the newborn infant. Cognition 121, 127–132 (2011).

22. J. Fiser, R. N. Aslin, Statistical learning of new visual feature combinations by infants. Proc. Natl. Acad. Sci. 99, 15822–15826 (2002).

23. A. Fló, et al., Newborns are sensitive to multiple cues for word segmentation in continuous speech. Dev. Sci. 22, e12802 (2019).

24. A. Fló, L. Benjamin, M. Palu, G. Dehaene-Lambertz, Sleeping neonates track transitional probabilities in speech but only retain the first syllable of words. Sci. Rep. 12, 4391 (2022).

25. N. Z. Kirkham, J. A. Slemmer, S. P. Johnson, Visual statistical learning in infancy: evidence for a domain general learning mechanism. Cognition 83, B35–B42 (2002).

26. J. R. Saffran, E. K. Johnson, R. N. Aslin, E. L. Newport, Statistical learning of tone sequences by human infants and adults. Cognition 70, 27–52 (1999).

27. C. Xu, et al., Statistical learning in patients in the minimally conscious state. Cereb. Cortex 33, 2507–2516 (2022).

28. A. Fló, L. Benjamin, M. Palu, G. Dehaene-Lambertz, Statistical learning beyond words in human neonates. eLife 13, RP101802 (2025).

29. E. S. Isbilen, S. M. McCauley, E. Kidd, M. H. Christiansen, Statistically Induced Chunking Recall: A Memory-Based Approach to Statistical Learning. Cogn. Sci. 44, e12848 (2020).

30. A. E. Milne, C. I. Petkov, B. Wilson, Auditory and Visual Sequence Learning in Humans and Monkeys using an Artificial Grammar Learning Paradigm. Neuroscience 389, 104–117 (2018).

31. M. Buiatti, M. Peña, G. Dehaene-Lambertz, Investigating the neural correlates of continuous speech computation with frequency-tagged neuroelectric responses. NeuroImage 44, 509– 519 (2009).

32. A. C. Schapiro, N. B. Turk-Browne, K. A. Norman, M.M. Botvinick, Statistical learning of temporal community structure in the hippocampus. Hippocampus 26, 3–8 (2016).

33. A. E. Kahn, E. A. Karuza, J. M. Vettel, D. S. Bassett, Network constraints on learnability of probabilistic motor sequences. Nat. Hum. Behav. 2, 936–947 (2018).

34. A. E. Kahn, D. S. Bassett, N. D. Daw, Trial-by-trial learning of successor representations in human behavior. PLOS Comput. Biol. 21, e1013696 (2025).

35. E. A. Karuza, A. E. Kahn, S. L. Thompson-Schill, D. S. Bassett, Process reveals structure: How a network is traversed mediates expectations about its architecture. Sci. Rep. 7, 1–9 (2017).

36. E. A. Karuza, A. E. Kahn, D. S. Bassett, Human Sensitivity to Community Structure Is Robust to Topological Variation. Complexity 2019, 8379321 (2019).

37. J. L. Elman, Finding Structure in Time. Cogn. Sci. 14, 179– 211 (1990).

38. G. Bellec, et al., A solution to the learning dilemma for recurrent networks of spiking neurons. Nat. Commun. 11, 3625 (2020).

39. X. Ren, H. Zhang, H. Luo, Dynamic emergence of relational structure network in human brains. Prog. Neurobiol. 219, 102373 (2022).

40. J. C. R. Whittington, et al., The Tolman-Eichenbaum Machine: Unifying Space and Relational Memory through Generalization in the Hippocampal Formation. Cell 183, 1249–1263.e23 (2020).

41. G. G. Ambrus, et al., When less is more: Enhanced statistical learning of non-adjacent dependencies after disruption of bilateral DLPFC. J. Mem. Lang. 114, 104144 (2020).

42. L. J. Batterink, D. Choi, Optimizing steady-state responses to index statistical learning: Response to Benjamin and colleagues. Cortex 142, 379–388 (2021).

43. L. J. Batterink, K. A. Paller, Statistical learning of speech regularities can occur outside the focus of attention. Cortex 115, 56–71 (2019).

44. L. Benjamin, G. Dehaene-Lambertz, A. Fló, Remarks on the analysis of steady-state responses: Spurious artifacts introduced by overlapping epochs. Cortex 142, 370–378 (2021).

45. H. Liu, T. A. Forest, K. Duncan, A. S. Finn, What sticks after statistical learning: The persistence of implicit versus explicit memory traces. Cognition 236, 105439 (2023).

46. J. M. Toro, S. Sinnett, S. Soto-Faraco, Speech segmentation by statistical learning depends on attention. Cognition 97, B25–B34 (2005).

47. I. Momennejad, Learning Structures: Predictive Representations, Replay, and Generalization. Curr. Opin. Behav. Sci. 32, 155–166 (2020).

48. K. L. Stachenfeld, M. M. Botvinick, S. J. Gershman, The hippocampus as a predictive map. Nat. Neurosci. 20, 1643– 1653 (2017).

49. E. D. Thiessen, A. T. Kronstein, D. G. Hufnagle, The extraction and integration framework: A two-process account of statistical learning. Psychol. Bull. 139, 792–814 (2013).

50. A. Christophe, E. Dupoux, J. Bertoncini, J. Mehler, Do infants perceive word boundaries? An empirical study of the bootstrapping of lexical acquisition. J. Acoust. Soc. Am. 95, 1570–1580 (1994).

51. M. Shukla, M. Nespor, J. Mehler, An interaction between prosody and statistics in the segmentation of fluent speech. Cognit. Psychol. 54, 1–32 (2007).

52. A. D. Endress, J. Mehler, The surprising power of statistical learning: When fragment knowledge leads to false memories of unheard words. J. Mem. Lang. 60, 351–367 (2009).

53. F. Al Roumi, S. Marti, L. Wang, M. Amalric, S. Dehaene, Mental compression of spatial sequences in human working memory using numerical and geometrical primitives. Neuron 109, 2627–2639.e4 (2021).

54. F. Al Roumi, S. Planton, L. Wang, S. Dehaene, Brain-imaging evidence for compression of binary sound sequences in human memory. eLife 12, e84376 (2023).

55. S. Dehaene, F. Al Roumi, Y. Lakretz, S. Planton, M. Sablé-Meyer, Symbols and mental programs: a hypothesis about human singularity. Trends Cogn. Sci. 26, 751–766 (2022).

56. S. Planton, et al., A theory of memory for binary sequences: Evidence for a mental compression algorithm in humans. PLOS Comput. Biol. 17, e1008598 (2021).

57. J. Quilty-Dunn, N. Porot, E. Mandelbaum, The best game in town: The reemergence of the language-of-thought hypothesis across the cognitive sciences. Behav. Brain Sci. 46, e261 (2022).

58. G. F. Marcus, K. J. Fernandes, S. P. Johnson, Infant Rule Learning Facilitated by Speech. Psychol. Sci. 18, 387–391 (2007).

59. F. Bulgarelli, D. J. Weiss, Anchors aweigh: The impact of overlearning on entrenchment effects in statistical learning. J. Exp. Psychol. Learn. Mem. Cogn. 42, 1621–1631 (2016).

60. A. L. Gebhart, R. N. Aslin, E. L. Newport, Changing Structures in Midstream: Learning Along the Statistical Garden Path. Cogn. Sci. 33, 1087–1116 (2009).

61. J.A. Jungé, B. J. Scholl, M. M. Chun, How is spatial context learning integrated over signal versus noise? A primacy effect in contextual cueing. Vis. Cogn. 15, 1–11 (2007).

62. J. M. Toro, J. B. Trobalón, Statistical computations over a speech stream in a rodent. Percept. Amp Psychophys. 67, 867– 875 (2005).

63. C. Santolin, O. Rosa-Salva, L. Regolin, G. Vallortigara, Generalization of visual regularities in newly hatched chicks (Gallus gallus). Anim. Cogn. 19, 1007–1017 (2016).

64. L. S. James, H. Sun, K. Wada, J. T. Sakata, Statistical learning for vocal sequence acquisition in a songbird. Sci. Rep. 10, 1–18 (2020).

65. O. Menyhart, O. Kolodny, M. H. Goldstein, T. J. DeVoogd, S. Edelman, Juvenile zebra finches learn the underlying structural regularities of their fathers’ song. Front. Psychol. 6 (2015).

66. A. Avarguès-Weber, et al., Different mechanisms underlie implicit visual statistical learning in honey bees and humans. Proc. Natl. Acad. Sci. 117, 25923–25934 (2020).

67. C. M. Conway, M. H. Christiansen, Modality-Constrained Statistical Learning of Tactile, Visual, and Auditory Sequences. J. Exp. Psychol. Learn. Mem. Cogn. 31, 24–39 (2005).

68. E. K. Nicholls, N. Hempel de Ibarra, Bees associate colour cues with differences in pollen rewards. J. Exp. Biol. 217, 2783– 2788 (2014).

69. M. D. Hauser, E. L. Newport, R. N. Aslin, Segmentation of the speech stream in a non-human primate: statistical learning in cotton-top tamarins. Cognition 78, B53–B64 (2001).

70. M. Boros, et al., Neural processes underlying statistical learning for speech segmentation in dogs. Curr. Biol. 31, 5512–5521.e5 (2021).

71. M. Takahasi, H. Yamada, K. Okanoya, Statistical and Prosodic Cues for Song Segmentation Learning by Bengalese Finches (Lonchura striata var. domestica). Ethology 116, 481– 489 (2010).

72. P. Perruchet, S. Desaulty, A role for backward transitional probabilities in word segmentation? Mem. Amp Cogn. 36, 1299– 1305 (2008).

73. B. Pelucchi, J. F. Hay, J. R. Saffran, Learning in reverse: Eight-month-old infants track backward transitional probabilities. Cognition 113, 244–247 (2009).

74. K. Tummeltshammer, D. Amso, R. M. French, N. Z. Kirkham, Across space and time: infants learn from backward and forward visual statistics. Dev. Sci. 20, e12474 (2016).

75. A. D. Endress, Learning melodies from non-adjacent tones. Acta Psychol. (Amst.) 135, 182–190 (2010).

76. E. L. Newport, M. D. Hauser, G. Spaepen, R. N. Aslin, Learning at a distance II. Statistical learning of non-adjacent dependencies in a non-human primate. Cognit. Psychol. 49, 85– 117 (2004).

77. R. Sonnweber, A. Ravignani, W. T. Fitch, Non-adjacent visual dependency learning in chimpanzees. Anim. Cogn. 18, 733–745 (2015).

78. E. Kakaei, S. Aleshin, J. Braun, Visual object recognition is facilitated by temporal community structure. Learn. Amp Mem. 28, 148–152 (2021).

